# *Trypanosoma cruzi* VDU deubiquitinase mediates surface protein trafficking and infectivity

**DOI:** 10.1101/2023.04.17.537123

**Authors:** Normanda Souza-Melo, Nathalia Brito Silva, Amaranta Muniz Malvezzi, Gregory Pedroso dos Santos, Laura Maria Alcântara, Carolina de Lima Alcantara, Narcisa Leal da Cunha e Silva, Martin Zoltner, Mark C. Field, Sergio Schenkman

## Abstract

Ubiquitylation is a post-translational modification promoting protein degradation. Within the endomembrane system ubiquitylation marks proteins for lysosomal processing. Deubiquitinases (DUBs) cleave ubiquitin from modified proteins and, in the case of surface proteins, prevent lysosomal targeting and thus play a major role in controlling turnover. In *Trypanosoma cruzi,* the etiological agent of Chagas disease, acquisition of nutrients in parasites proliferating within the blood meal of insect vectors occurs via the cytostome, a unique structure connected to a long tubular cytopharynx. Contents are delivered to late endosomes and accumulate in reservosomes, equivalent to lysosomes. When starved, *T. cruzi* differentiates into mammalian infective trypomastigotes, which are cell cycle arrested. Here we asked what roles ubiquitylation plays in this unique endocytic process by interrogation of the *T. cruzi* ortholog of VDU (von Hippel-Lindau-interacting deubiquitylating enzyme)/USP33 (ubiquitin-specific protease). We found that TcVDU expression level inversely correlated with transferrin endocytosis, and that overexpression led to a longer retention of the endocytic cargo near the cytostome. TcVDU itself was found enriched in the anterior region of the parasite, in proximity to endocytic cargo. Most importantly, TcVDU overexpression reduced parasite invasion capacity and led to increased release of *trans*-sialidase. These alterations in the abundance of multiple surface proteins in TcVDU mutants indicate a key role of TcVDU in modulating the *T. cruzi* surface by affecting the endosomal traffic and consequently the host-parasite interface.

**Author summary:** *Trypanosoma cruzi* is the cause of Chagas disease that affects large populations of Central and South America. Disease is spreading to other continents due to poor control of blood transfusion from donors with chronic and undiagnosed infection, migration and drug treatment with limited performance and undesired side effects. Proliferating forms of *T. cruzi* acquire nutrients through the cytostome, a cell surface opening connected to a long tubular cytopharynx. Internalized material is endocytosed in the cytopharynx. This is distinct from other members of the Trypanosomatids, which endocytose material exclusively through the flagellar pocket. By using CRISPR gene editing and overexpression we demonstrated that a conserved deubiquitinase (VDU) is a key control element for traffic from the cytopharynx to endosomal compartments in *T. cruzi*. Changed levels of VDU modify endosomal traffic and largely impact the parasite surface, causing changes in infectivity.

## Introduction

Ubiquitination is a reversible post-translational modification forming an isopeptide bond between ubiquitin and a protein client, facilitating regulation of many functions, including degradation, targeting and cell cycle control [1]. Deubiquitinase proteases (DUBs) remove ubiquitin and are key regulators of these process being involved in controlling protein degradation, endocytosis, DNA repair and membrane protein traffic [2, 3]. A cohort of about a dozen DUBs are present in trypanosome genomes [4], several of which are conserved across eukaryotes [5].

*Trypanosoma cruzi* is the causative agent of Chagas disease, which affects about seven million people worldwide [6, 7]. It is a flagellated protist parasite with a complex life cycle [8–10]. The plasma membrane is dominated by many distinct glycosylphosphatidylinositol (GPI) anchored glycoconjugates, which change throughout the different parasite life cycle stages [11–13], which are mainly mediated by differential gene expression [14, 15]. In addition, surface component abundance is controlled by the membrane trafficking system [16]. For example, the epimastigote form, which replicate in the gut of the insect vector, has a high capacity to accumulate material from a blood meal into endocytic vesicles known as reservosomes (equivalent to lysosomes) [17–21], while the surface is predominantly composed of glycosylphospholipids [13, 22]. Amastigotes, the life stage replicating inside mammalian cells, also perform endocytosis to acquire host components [17, 23] and express specific glycoproteins such as amastins [24] and mannose enriched glycoproteins [25]. In contrast, non-proliferative infective trypomastigotes have low endocytic capacity [26, 27] and express in the surface and secrete large amounts of GPI-anchored mucin-like glycoproteins [28], along with specific members of the *trans*-sialidase gene family [29]. These include active *trans*-sialidases and are important and modulate several host interactions [15, 29–31]. Therefore, alterations between life cycle stages occurs in parallel to changes of the membrane trafficking system, but it remains unclear how this occurs, and is unlikely simply a result of altered gene expression.

In *Trypanosoma brucei,* a related protozoan that causes African trypanosomiasis, intracellular traffic of several surface proteins depends on coupled reactions of ubiquitination and deubiquitination [32], which includes the invariant membrane proteins ISG65 and ISG75. ISG75 is the receptor for the trypanocide suramin as well as other structurally related compounds [33]. ISG65 servers as a complement receptor with a role in complement inactivation [34, 35]. Silencing trypanosome orthologs of the highly conserved DUBs USP7 and VDU render resistance to suramin by preventing delivery to the lysosome and by decreasing levels of ISG75 [36]. Whilst the mode of action of suramin appears to be highly complex [36], it is clear that VDU1 and USP7 are important mediators of surface protein turnover. As orthologs of both USP7 and VDU1 are encoded in the genome of *T. cruzi,* we hypothesized that they play a similar role in this related parasite, potentially controlling expression, secretion, or turnover of surface proteins in the different life cycle stages of *T. cruzi*. In mammalian cells VDU1 is involved in the control of the von Hippel-Lindau E3 ligase and its dysregulation can lead to multiple forms of tumorigenesis [37], hypothyroidism and additional morbidities [38]. VDU1 also acts on Robo1 amongst its substrates, which may be the closest parallel to functions in trypanosomes, as Robo1 is a *trans*-membrane domain receptor for the cell migration ligand Slit [39].

As a first approach to understanding the roles of DUBs in life cycle progression and modulation of the parasite surface proteome, we performed overexpression and knockout of TcVDU and investigated the impact on endocytosis and differentiation. We found that TcVDU is a major modulator of the surface proteome and endocytic activity with a significant impact on the parasite life cycle.

## Results

### Sequence analysis of TcVDU

The ubiquitin carboxyl-terminal hydrolase (UCH) in *T. cruzi*, TcVDU (BCY84_01319) is orthologous and syntenic with *T. brucei* and *Leishmania mexicana*, with 44% and 31.7% predicted amino acid identity, respectively. The gene encodes a protein of 761 amino acids with a predicted molecular weight of 87 kDa. By BLAST analysis with TcVDU CDS using Interpro, we found 14 similar human DUB proteins. A phylogenetic tree using MrBayes is shown in Figure 1A and indicates that not only *T. cruzi*, but also *T. brucei* and *L. mexicana* are closely related to human USP38, USP47, USP33 and USP20 DUBs. Domain analysis indicates that TcVDU has a comparable domain arrangement to USP33 and USP20, as detected by Pfam analysis (Figure 1B). The strongest resemblance with USP33 and USP20, a recent duplication, was due to the presence of the conserved catalytic triad in TcVDU corresponding to Cys49, His569, and Asp589 residues (Figure 1C) [40, 41]. These residues are also present in the *T. brucei* (Tb927.11.12240) and *L. mexicana* VDU orthologs (DUB3, LmxM.29.1200). Interestingly, trypanosome VDUs lack the zinc finger domain, involved in the recognition of the C-terminal of ubiquitin by USP33 [42], and the DUSP domain, typically spanning 120 amino acid residues, and involved in USP33 interactions [43], suggesting functional differences in these parasites.

**Figure 1:**
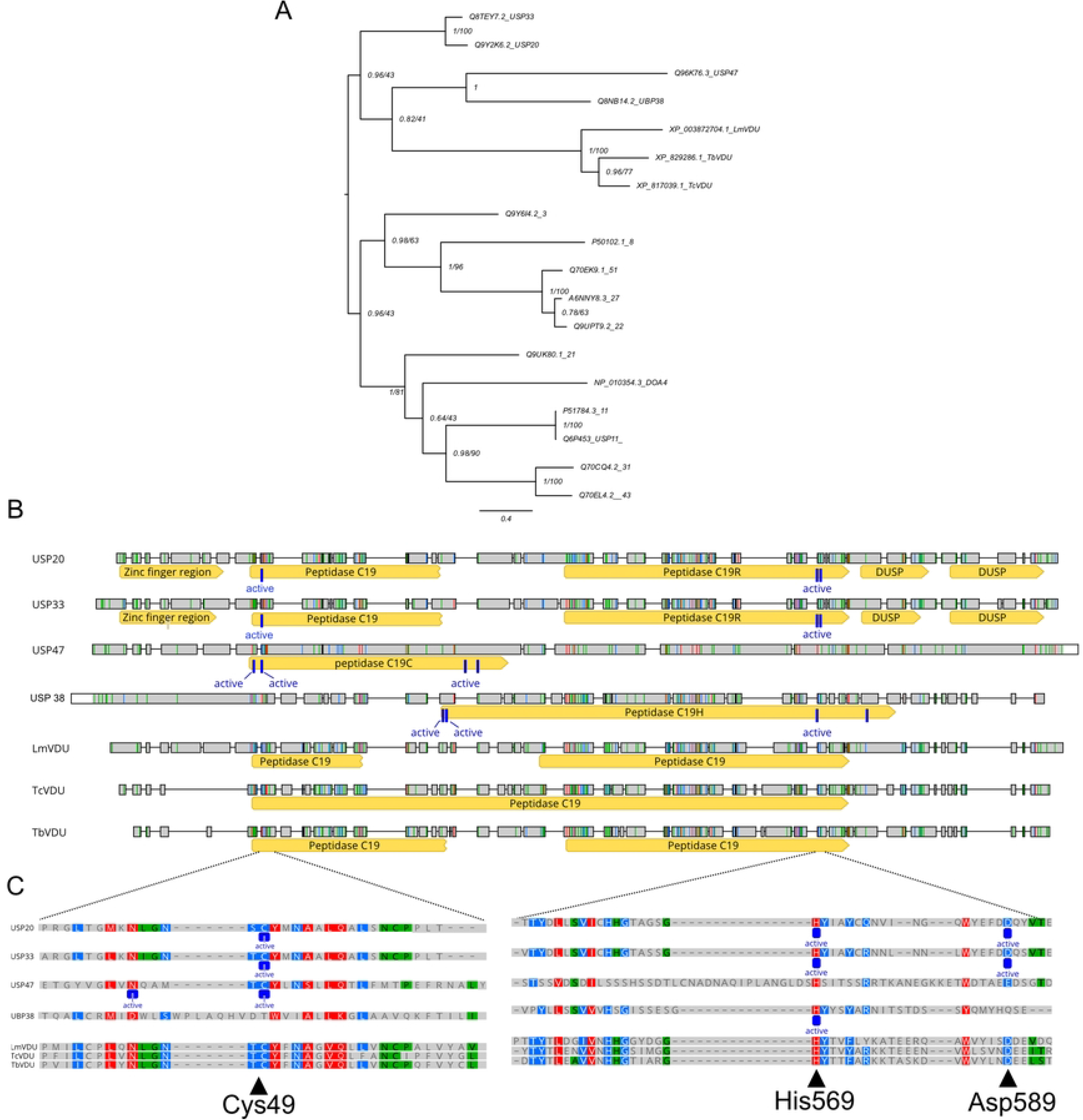
Ubiquitin Carboxyl-terminal Hydrolase (UCH) sequences of kinetoplastids are homologs to the human UCH sequences. A) Phylogenetic tree of human DUBs similar to the kinetoplastid VDU proteins generated by PhyML (100 nonparametric and standard bootstraps and MrBayes settings with four chains run with a sampling frequency of 500 to an average standard deviation of splits frequencies <0.01, indicating convergence. The numbers indicate 100 Bootstraps out of 1.0 to test branches. The VDU sequences of T. cruzi (TcBCY84_01319), T. brucei (Tb927.11.12240), and L. mexicana (LmxM.09.0240) were from https://tritrypdb.org/tritrypdb/). Homo sapiens deubiquitinases were respectively for USP33, USP20, USP47, USP38, USP22, USP27, USP8, USP3, USP43, USP31, UCTH21, USP43, USP11: Q8TEY7.2, Q9Y2K6.2, Q96K76.3, Q8NB14.2, Q9UPT9.2, A6NNY8.3, Q70EK9.1, P50102.1, Q9Y6I4.2, Q9UK80.1, Q70EL4.2, Q70CQ4.2, P51784.3 (https://www.uniprot.org/). B) Schematic representation of the alignment of VDU proteins of T. cruzi, T. brucei, and L. mexicana with the closest hits of human deubiquitinases identified by using InterPro and redrawn with the Geneious software [118]. Similar sequences are indicated by colored boxes (red > 80%, blue > 60%, green > 40% and gray for other residues). Homo sapiens deubiquitinases were respectively for USP33, USP20, USP47, USP38, USP22, USP51: Q8TEY7.2, Q9Y2K6.2, Q96K76.3, Q8NB14.2, Q9UPT9.2 (https://www.uniprot.org/). The bars indicate the Interpro protein domains and the dark blue lines the amino acids of the active site of the enzymes. DUSP refers to a domain conserved in deubiquitinases. C). Sequence alignment detail of TcVDU active site regions; the positions of active site residues Cys49, His569, and Asp589 are indicated.

### Impact of VDU overexpression and knockout

As *T. brucei* VDU is involved in controlling membrane traffic we decided to obtain *T. cruzi* parasites with overexpressed and depleted VDU. For overexpression, we generated parasite lines by stable acquisition of a plasmid with a strong ribosomal promoter [44, 45] containing the VDU gene fused to GFP, termed VDU^GFP^ (Figure 2A). The parasite line displayed GFP fluorescence by flow cytometry (Figure 2B), a protein band with an apparent molecular weight of 120 to 135 kDa consistent with the predicted size of the fusion protein (120 kDa), which was recognized by anti-GFP antibodies (Figure 2C and Figure S1A). The observed shift in molecular weight may be due to post-translational modifications such as N-glycosylation, affecting migration in SDS-PAGE gels [46]. VDU^GFP^ was enriched near the kinetoplast, a region corresponding to the flagellar pocket or cytostome/cytopharynx entrance (Figure 2D). In addition, TbVDU showed cytoplasmic localization around the nucleus and defined regions in the anterior region of the parasite. A similar distribution was present in intracellular amastigotes (Figure S2). Endogenous tagging with an N-terminal V5 sequence carried out to avoid overexpression and disruption of potential regulatory at the 3’UTR [47], led to a similar cellular distribution of the fusion protein and there was some enrichment in the region close to the flagellar pocket (Figure S3). In contrast, GFP expressed alone is present throughout the parasite, indicating that accumulation near the flagellar pocket was not a result of the overexpression (Figure 2D). In parallel, we generated parasites that overexpressed USP7, previously found to participate in endocytic trafficking in *T. brucei* [32], at similar levels of VDU as observed by Western blotting and flow cytometry (Figures S1A-C).

**Figure 2:**
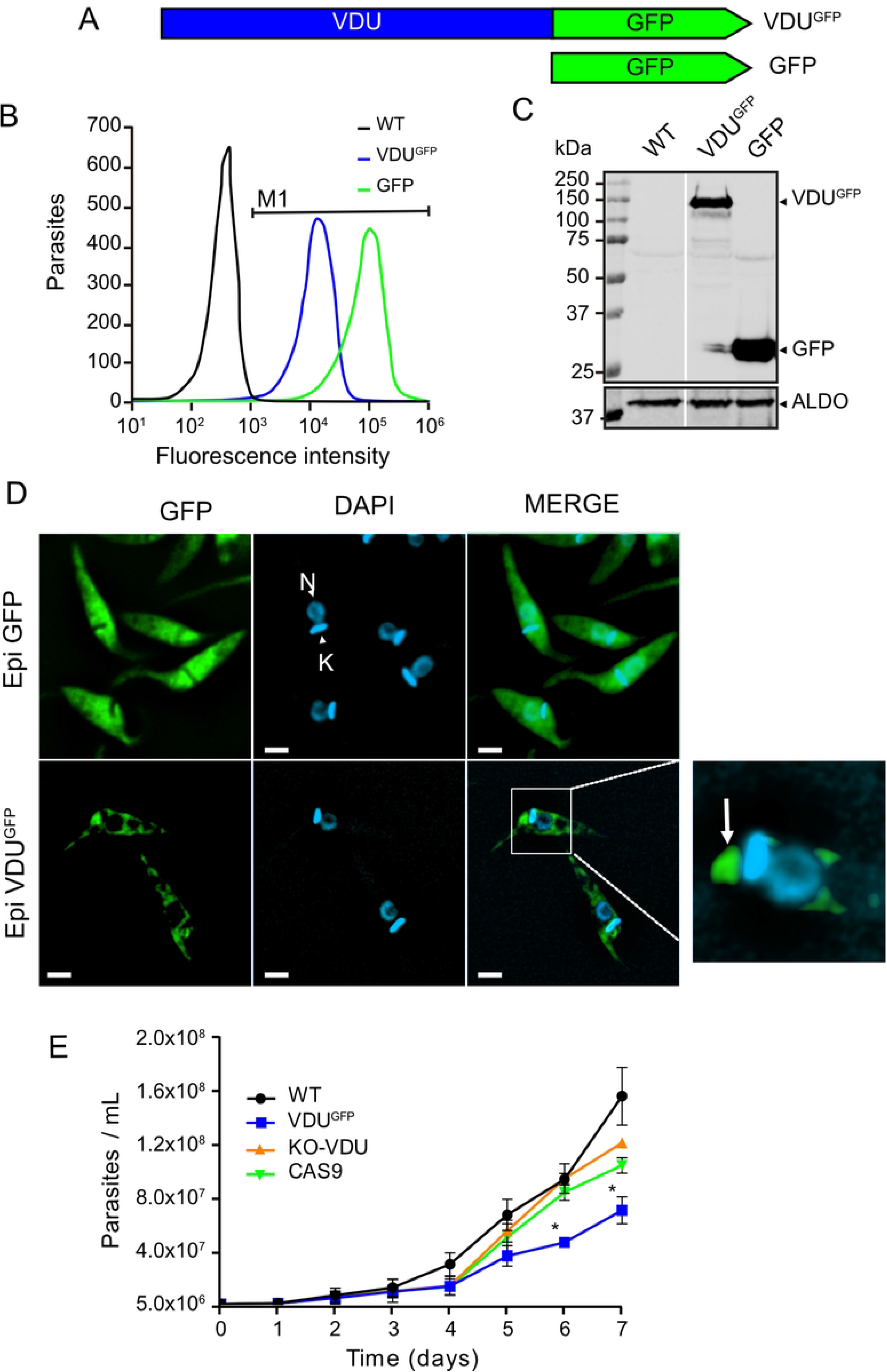
Expression and localization of TcVDU-GFP in T. cruzi epimastigote. A) Scheme of TcVDU fused to GFP (VDU^GFP^) and the plasmid expressing GFP alone. B) Representative of flow cytometry of several observations of live epimastigotes selected for G-418 resistance after transformation with plasmids carrying VDU^GFP^ (blue) or GFP (green) in comparison with wild-type (WT) *T. cruzi* Dm28c strain (black). C) Western blot of selected cell lines decorated with the anti-GFP antibody, representative of two experiments. The molecular mass of standards is indicated in kDa, and arrowheads highlight the migrating position of VDU^GFP^ (114 kDa), GFP (27 kDa), and aldolase (ALDO – 41 kDa) controls. D) Fluorescence microscopy of epimastigotes expressing GFP alone (Epi-GFP), or VDU^GFP^. The panels show the GFP and DAPI and the merged fluorescence. Bars correspond to 2 µm. N: nucleus. K: Kinetoplast. (E) Growth curves of WT, Cas9, VDU^GFP^ and KO-VDU epimastigotes. Each experimental point represents the mean ±SD of triplicate experiments. Asterisks indicate p ≤ 0.001 when compared to wild type cells, using two-way ANOVA test.

To explore how loss of TcVDU expression alters endocytic processes, we additionally generated knockout parasites using CRISPR-Cas9 by replacing the N-terminal portion of the gene with the BSD resistance gene (Figure S4A), as previously described [48, 49]. After selection with BSD correct insertion was verified by PCR. DNA from wild-type and Cas9GFP parasites generated a fragment of 958 bp whereas the KO-VDU presented a 1173 bp (Figure S4B). Oligonucleotides P3 and P4, which recognize the BSD coding region, confirmed insertion of the resistance gene in the KO line (399 bp, Figure S4C). PCR further confirmed the absence of the N-terminal VDU portion (Figure. S4D). In addition, we found a very large reduction (>50X) of TcVDU mRNA qPCR, but not complete depletion, probably due to the presence of incomplete selection (Figure S4E). More importantly, a similar reduction in expression was observed with primers annealing upstream and downstream the Cas9 cleavage site, rendering it unlike that a truncated TbVDU protein is expressed along the inserted BSD resistance protein. The absence of VDU expression did not affect parasite proliferation *in vitro* as compared to wild-type or Cas9 parasites (Fig. 2E), but replication was significantly reduced by overexpressing VDU^GFP^. As epimastigote forms obtain some of their nutrients by a continuous endocytic process [50], we examined the endocytic activity in this parasite stage.

### VDU^GFP^ overexpression decreased while VDU knockout increased the incorporation of transferrin by *T. cruzi*

In eukaryotic cells, endocytosis is in part regulated by ubiquitination. Many surface receptors are ubiquitinated, which signals transport to secondary endosomes and fusion with lysosomes to be degraded. Deubiquitination allows recycling to the surface and hence increases abundance [51, 52]. Epimastigote forms can continuously obtain nutrients through the cytostome-cytopharynx [19, 53], which disassemble upon differentiation into non-proliferative and infective forms [27, 54]. Thus, we tested the endocytic capacity of epimastigotes overexpressing or depleted of VDU for uptake of fluorescently labeled transferrin (TF^633^). Initially, parasites were incubated for 15 min in medium without serum to allow internalization of unlabeled transferrin, and then incubated with transferrin-Alexa 633 (TF^633^) for 30 min. Flow cytometry revealed that TF^633^ uptake was decreased in VDU^GFP^ parasites, whereas an increased uptake was observed for the KO-VDU line (Figure 3A). These differences were not attributed to variations in the cell cycle, as all cell lines presented similar amounts of cells in G1, S and G2 phase (Figure 3B), which could affect endocytosis [55]. Quantitative analysis, performed in quadruplicate experiments, showed that the number of labeled parasites, or the total incorporated fluorescence in parasites decreased in VDU^GFP^ overexpressors compared with wild-type or Cas9 parasites (Figure 3C and D). In contrast, the KO-VDU showed a significative increase in TF^633^ incorporation. Individual VDU^GFP^ cells observed by microscopy had lower fluorescence compared to the Cas9 and KO-VDU, with the latter exhibiting brighter signals than the other cell lines (Figure 3E-F), confirming the flow cytometry. Furthermore, overexpression of USP7, a second DUB involved in endocytic protein trafficking in *T. brucei* [32] at a similar level to VDU did not inhibit transferrin endocytosis in *T. cruzi* epimastigotes (Figure S5), suggesting the effect of VDU overexpression is specific to this DUB [32]. Decreased endocytosis in VDU overexpressors line can explain its reduced proliferation rate.

**Figure 3:**
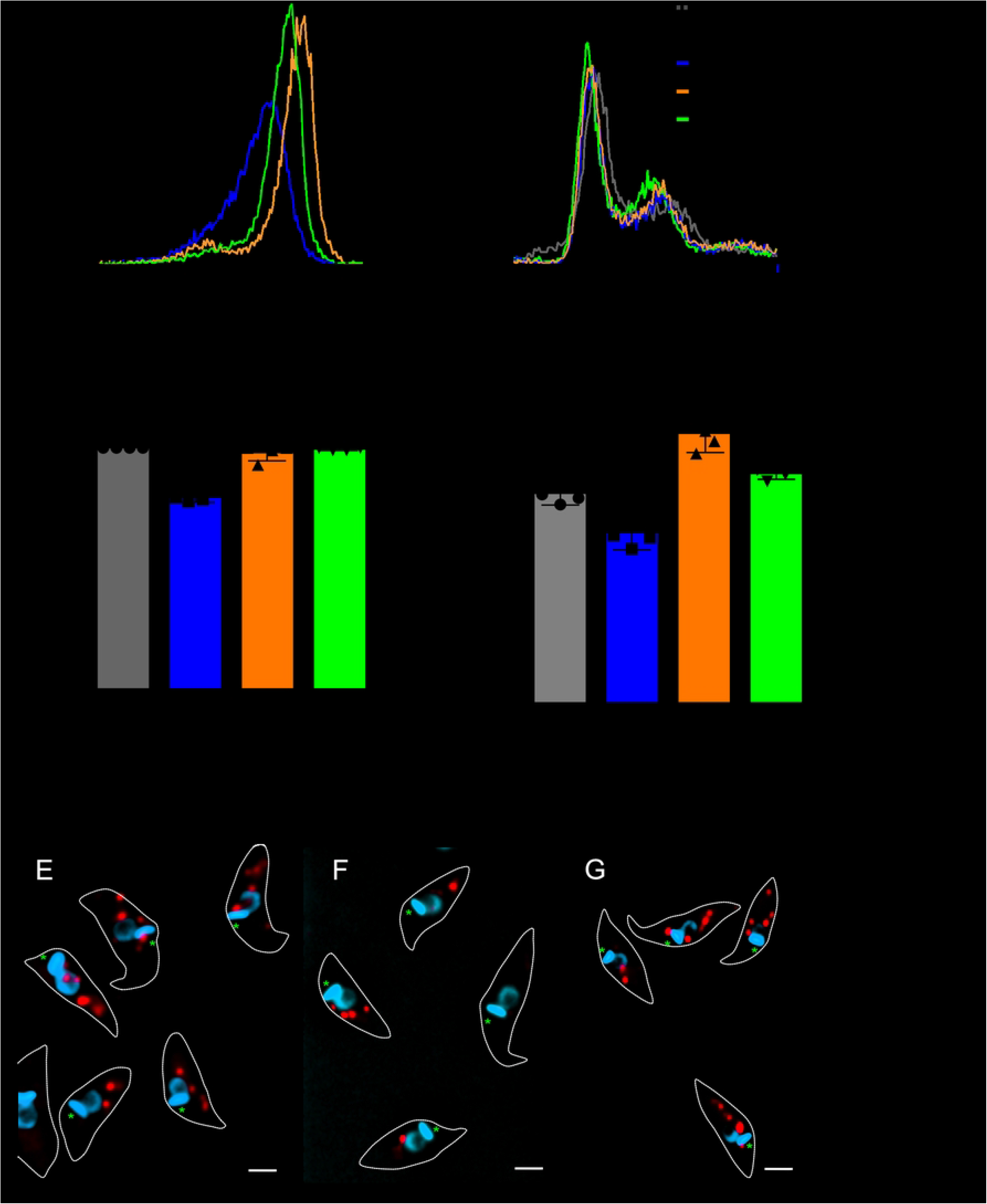
Endocytosis of TF^633^ in epimastigote is decreased in the VDU^GFP^ overexpressor and increased in the KO-VDU. (A) Representative flow cytometry histograms of epimastigotes of the indicated lines incubated with TF633 for 30 minutes in RPMI-164 medium, FBS-free at 28 °C. As a negative control, wild-type (WT) parasites were incubated without labeled transferrin (dashed line). (B) Cell cycle analysis of the same lines made after labeling with 7-AAD. Quantitative analysis of TF633 endocytosing cells measured as the number of fluorescent parasites calculated above the fluorescence intensity of 1000, which correspond to WT parasite fluorescence without TF^633^ () (C) and the total incorporation of TF^633^ calculated as the median fluorescence subtracted from the median of parasites incubated without TF^633^ (D), both analyzed upon 30 min incubation. Each graphics shows the mean of 4 biological replicates with mean ± SD. Asterisks represent the significance relative to WT cells with p ≤ 0.0001 (****), p ≤ 0.001 (***), and p ≤ 0.01 (**) with respect to wild type. Hash symbols are p ≤ 0.001 (# # #) and p ≤ 0.01 (# #) in relation to Cas9. A one-way ANOVA test was used to assess the statistical difference between samples. (E, F and G) show typical fluorescence microscopy images obtained from the experiment above. Blue corresponds to DAPI staining of the nucleus and kinetoplast, red to TF^633^. The dashed lines indicate the parasite contour and the green asterisk the approximate position of the flagellar pocket.

Next, we evaluated the kinetics of TF^633^ trafficking through the endocytic pathway. *T. cruzi* epimastigotes obtain nutrients via the cytostome-cytopharynx. complex, localized in the anterior region of the parasite, from which the flagellum protrudes [56, 57]. The anterior region is easily assigned as it is opposite to the nucleus (N) with respect to the kinetoplast (K), an elongated DNA-containing structure strongly labeled with 4.6-diaminodino-2-phenylidole (DAPI). Over time, endocytic cargo migrates from the anterior (A) region, passing close to the nucleus (N), to the posterior end of the parasite (P), where endocytosed material accumulates in endosomes for storage or other uses, or degradation (reservosomes) (Figure 4A). The uptake of labeled TF^633^ was monitored by fluorescence microscopy. After 1 minute we observed very weak labeling in the anterior region in controls (WT and Cas9) and in KO-VDU parasites (Figure 4B). Most of the signal was in the N region at this time point. In contrast, we found a significantly higher number of parasites labeled at the A region in the VDU^GFP^ cells. After 10 min incubation, labeling was dominant at the P region in all cell lines except VDU^GFP^ (Figure 4C). Within 30 min, an evident accumulation was observed in the P region, but the VDU^GFP^ line retained significant levels in the M region (Figure 4D). Altogether, these results indicate a delay in the traffic of transferrin to the posterior part of the cells in cells overexpressing VDU^GFP^.

**Figure 4:**
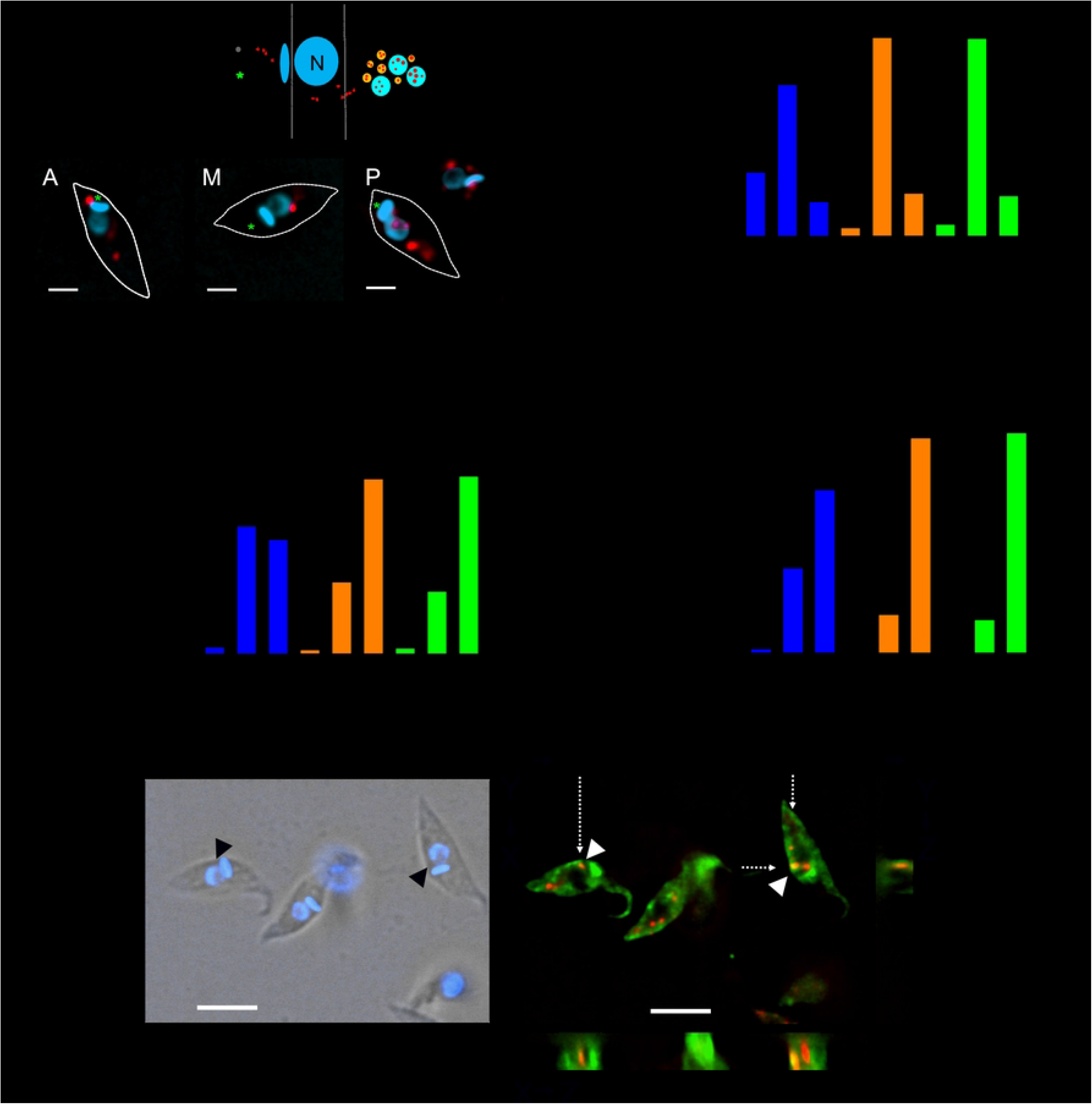
The VDU^GFP^ overexpressor shows a reduced rate of endocytosis of TF633. (A) Schematic representation of the dynamics of TF^633^ endocytosis and representative images of transferrin trafficking stages in *T. cruzi* cells obtaining from random fields (Bars = 2 mm). Epimastigotes were incubated with TF^633^ for 1, 10, or 30 min in low glucose DMEM medium, without FBS at 28°C. At the indicated time points, parasites were fixed with paraformaldehyde, the DNA was stained with DAPI, and samples were analyzed by fluorescence microscopy. The images show the DNA labeling in blue and TF^633^ in red. A) represents the pattern of transferrin in the anterior region and close to the kinetoplast; M: in the middle of the parasite corresponds to intermediate stage of accumulation; and P: accumulation in the posterior region, which correspond to the secondary endosomes. The number of each pattern was quantified in at least 100 hundred parasites after 1 (B), 10 (C), and 30 min (D). The bars depict the mean ± SD of 3 independent experiments with n = 100. The asterisks representing the significance of p ≤ 0.05 (*) in relation to WT, calculated using one-way ANOVA test. E) Endocytosis experiments were performed by incubation of parasites in RPMI medium with TF^633^ for 10 min at 28°C. The parasites were then fixed for 20 min with 4% p-formaldehyde in PBS, washed in PBS and attached to poly-L-lysine coated glass slides. The slides were then treated with DAPI and observed in the fluorescence microscopes. The images correspond to one Z-section generated after blind deconvolution, showing the VDU^GFP^ (green), TF^633^ (red), and DAPI (blue). White arrowheads indicate the superposition of TF633 and VDUGFP generate the images in XY, XZ and YZ axis. The left panel shows the DIC and DAPI superimposed images. The right panel shows the VDU^GFP^ and TF^633^ images using the 3 planar projections, as indicated. The arrowheads indicate the superimposition of the two fluorescent markers, also shown as black arrowheads in the top panel, pointing out that VDU is not broadly localized The dashed arrows show the position used to generate the Z-axis. Bar = 5 µm.

### Some TcVDU localizes close to *T. cruzi* endocytic structures

To further analyze at which point of the endocytic process TcVDU acts, VDU^GFP^ cells were monitored by microscopy after incubation with TF^633^. As shown in the Figure S6, we observed some endocytic structures with TF^633^ at locations also exhibiting VDU^GFP^ fluorescence. The observed superposition was detected mainly in the M region of the cell body, in which a long stretch of green fluorescent is seen. The eventual proximity along the Z axis is shown in Figure 4E, particularly at positions in which the cargo is in a region of the cytopharynx. The Z-projection and three-dimensional analysis of this image (Supplementary video V1) indicates that TF^633^ labeling appears surrounded by TcVDU-GFP fluorescence. Notably, TcVDU tagged with a V5 epitope was also more concentrated in the region proximal to the flagellar pocket, excluding the possibility that TcVDU overexpression led to abnormal accumulation in this region. Taken together, these images suggest that TcVDU delays transfer of endocytic cargo to the M region of the cell.

Similar observations were made by TEM of endocytosed transferrin coupled to colloidal gold (Tf-Au). The TF-Au tracer was localized after 30 min incubation. As expected, we observed endocytic uptake of gold particles in all cell lines. The particles were detected in small and medium size vesicles, which are presumed to correspond to early and late endosomes (E) and large reservosomes (R), the latter with a denser lumen and electron-lucent inclusions. In WT cells, both endosomes and reservosomes contained gold particles (Figure 5A and B). For the VDU^GFP^ mutant, we observed some parasites with an internalized TF-Au tracer pattern similar to WT. Some small vesicles with a high concentration of TF-Au were captured fusing with likely reservosomes (Figure 5C), but this was rarely observed. We also noticed accumulation of the endocytic tracer in endosomes localized to the posterior region of the cell and along the cytopharynx (Figure 5D). These findings are suggestive of a delay in cargo delivery to the endosomal network, fully consistent with the results with the TF^633^ tracer. KO-VDU cells showed similar patterns to WT but with a higher number of gold particles dispersed in the endosomes and reservosome lumen (Figure 5E and F). Interestingly, we were able to find large reservosomes in fusion, a structure rarely found in *T. cruzi* (Figure 5G).

**Figure 5:**
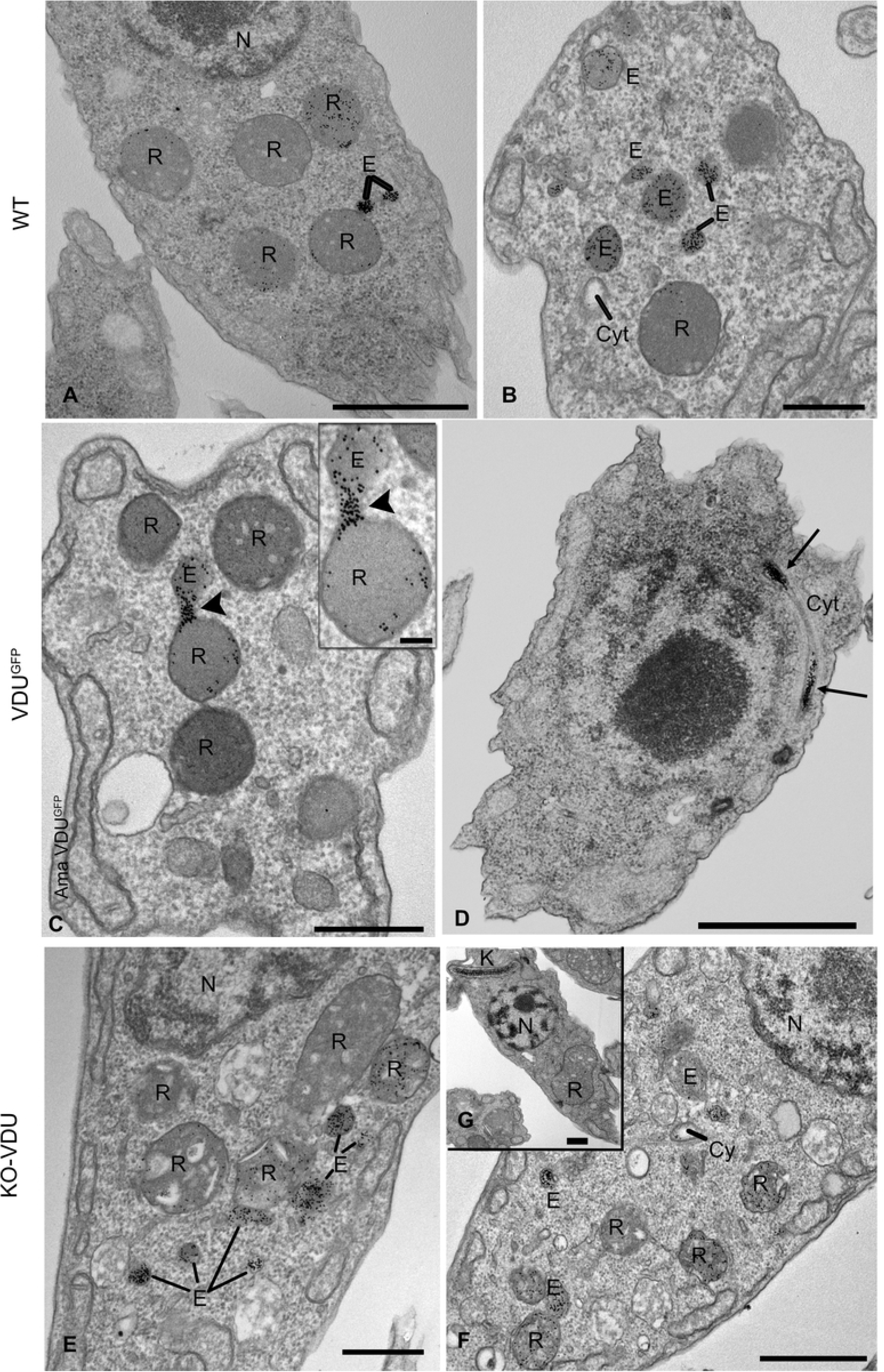
Endocytosis of transferrin-gold in VDU mutants. Epimastigotes were incubated with TF-Au for 30 min at 28°C. Samples were fixed and processed for TEM. A, B) in WT epimastigote, TF-Au could be found in typical endosomes (E) and large endosomes, considered reservosomes (R). C, D) In VDU^GFP^ overexpressors, TF-Au is present mainly in small endosomes and images (n = 20) show TF-Au being transferred to reservosomes is frequently observed (arrowhead in C, with higher magnification in the inset). We also detected TF-Au in the cytopharynx (Cyt) as indicated by the arrows in (D). E, F) KO-VDU mutants show endocytic pattern comparable to the WT, with presence of TF-Au inside endosomes and reservosomes, although reservosomes appear to contain more tracer when compared with WT. G) Shows a rare image of bilobate large reservosomes. N (nucleus), Cy (cytopharynx). Scale bars: A, D, E – 1 µm; B, C, F – 500 nm; inset bar in C and G – 100 nm.

We then performed three-dimensional reconstruction of parasite sections obtained by Focused Ion Beam-Scanning Electron Microscopy. We observed that KO-VDU cells presented larger reservosomes than WT cells by visual comparison (Figure 6A). For KO-VDU cells and parental cell line, reservosomes from entire cells in G1 and G2 stages of the cell cycle were reconstructed and number per cell and volume calculated. Quantitative analysis showed a significant increase in reservosome volume in G1 cells (Figure 6B), while differences were smaller and not statistically significant in G2 cells. No noticeable differences were observed in VDU^GFP^. Taken together, these findings support the notion that VDU is a negative regulator of the endocytic process in *T. cruzi*.

**Figure 6:**
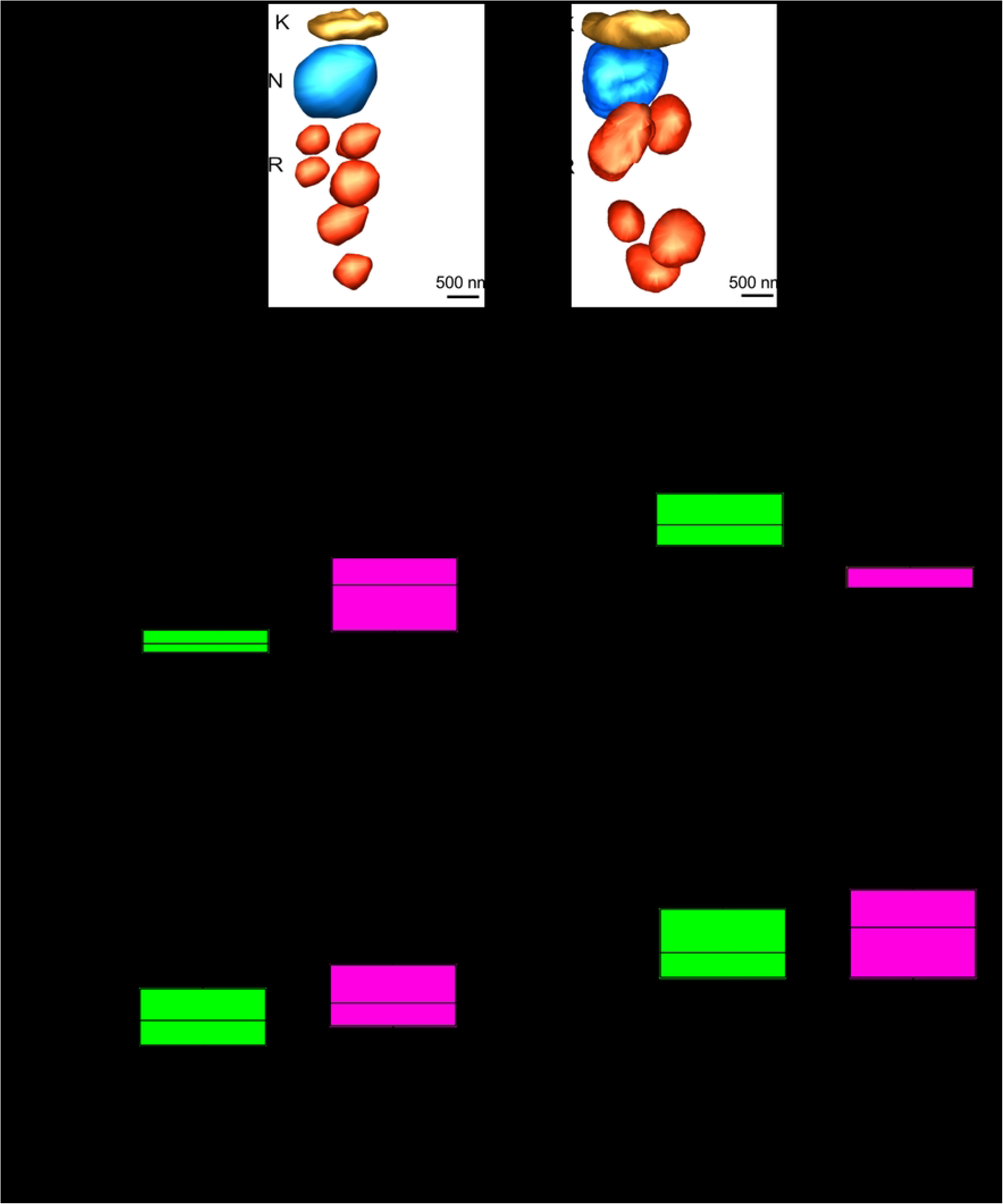
KO-VDU reservosomes are larger. (A) Three-dimensional reconstruction of WT (CAS9) and KO-VDU epimastigote in G1 phase of the cell cycle by FIB-SEM. The gold-colored structures correspond to the kinetoplast (K), blue to the nucleus (N) and red to reservosomes (R). Bars = 0.2 µm. Mean and standard deviation of the reservosome volume (B and D) and reservosome number/cell (C and E) of indicated cells in G1 and G2 calculated from the three-dimensional reconstructions (n = 6 and 5, respectively). Asterisk symbol indicates significant differences with p ≤ 0.05 in relation to Cas9 by using a one-way Anova test.

### TcVDU overexpression reduces parasite infectivity

Endocytic traffic is largely reduced upon differentiation of proliferative epimastigotes or amastigotes to infective non-proliferative trypomastigote forms [27]. Therefore, we investigated the consequences of perturbed endocytosis and membrane trafficking on differentiation of epimastigotes into metacyclic-trypomastigote forms and for tissue culture trypomastigote (TcT) infectivity. All cell lines generated trypomastigote forms (metacyclic-trypomastigote or TcT) that were competent to infect mammalian cells. However, both metacyclic-trypomastigote and TcT derived from VDU^GFP^ parasites appeared less infective compared to the KO-VDU and Cas9 parasites. To determine the molecular basis for this, we initially examined infections using similar numbers of TcT cells. After infection of U-2OS cells, we observed a significant decrease in the number of cells that presented intracellular parasites for the VDU^GFP^ mutant as compared to the KO-VDU and control parasites (Figure 7A). Next, using variable parasite numbers to obtain the same infectivity, we found that intracellular numbers of amastigotes were similar for both lineages 48 h post-infection (Figure 7B). Under the same conditions, all cell lines egressed from the infected mammalian cells without significant kinetic differences (Figure 7C). Importantly, the VDU^GFP^ was expressed in intracellular amastigotes mainly at the region of the cytostome/cytopharynx complex entrance and flagellar pocket region, but it was also visible as a more diffuse cytoplasmic signal along of the cellular body (Figure S2). These results suggest that changes promoted by an excess of VDU affect the invasion capacity of TcTs, thereby causing reduced infectivity, but not later events in the infection cycle.

**Figure 7:**
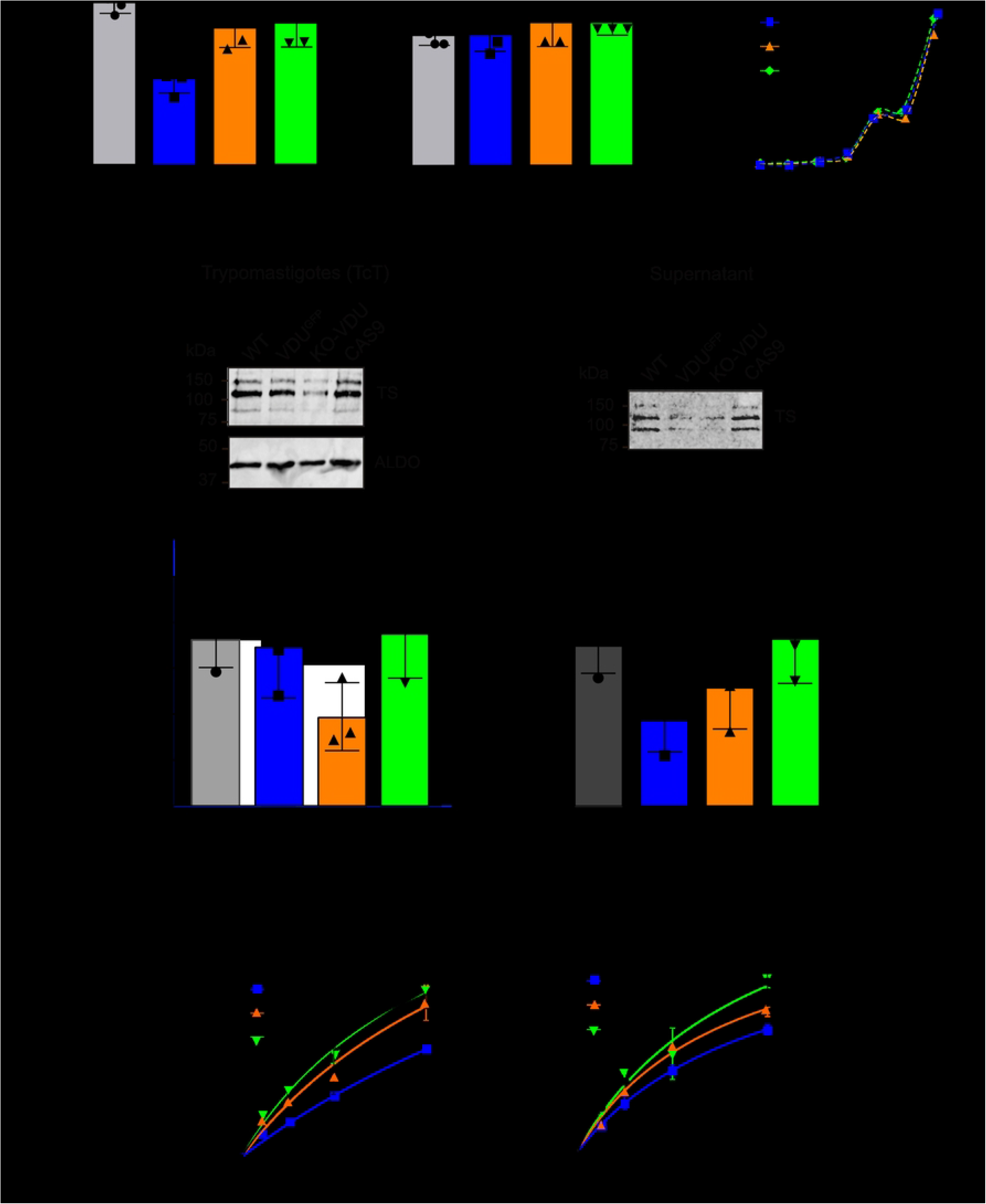
TcVDU^GFP^ overexpression decreases cell invasion and reduces trans-sialidase sialidase secretion in TcTs. U2OS cells were infected in the with TcT cells of the indicated lines at the same MOI (10:1), and infection ratio determined after 24 h, as described in methods (A). A new infection was carried out to obtain the same infectivity using proportional MOI, and the number of intracellular parasites (B) and trypomastigotes egress by direct counting using a Neubauer chamber (C). Each graphics represent a mean of 4 biological replicates with mean ±SD. (D-E) Representative immunoblots of TcT lysates, and the respective culture supernatant probed with mAb39 (anti-TS) (top) or anti-aldolase (bottom). (F-G) Graphical representations of TS/ALDO pixel intensity normalized to WT in each western blot (n=3). The graphics in (F, G) show the intensity of all TS bands in the parasites and in the supernatant relative to aldolase in the cells. Each value represents a mean of 2 technical replicates and 3 biological replicates with mean ± SD. The TS activity was measured with the indicated amounts of parasite lysates in a 30 min assay (H) and in the culture supernatant in a 2 h assay (I). The significances were calculated using two-way Anova test, with p ≤ 0.001 (***), p ≤ 0.01 (**) and p ≤ 0.05 (*).

### TcVDU overexpression reduces the levels and activity of *trans*-sialidase

*Trans*-sialidases (TS) are a major family of glycosylphosphatidylinositol anchored proteins predominantly expressed on the surface of trypomastigotes and are important virulence factors due to mediating interactions between the parasite and host cells [58–60]. The TS family includes members able to transfer sialic acid from host glycoconjugates to acceptors such as mucin-like glycoproteins also on the parasite surface and important to host cell invasion [61, 62]. In TcT, the TS subgroup contains unique 12 amino acid repeats originally identified as the shed acute phase antigen (SAPA) [12]. As the VDU^GFP^ mutant displayed a decrease in infection and both mutants exhibit altered endocytosis pathways, we examined the levels of TS containing SAPA in trypomastigote forms and culture supernatants. Western blots with mAb39, which only recognize TS with SAPA repeats [63], showed that the corresponding antigen was reduced in the KO-VDU mutant (Figure 7D and E). However, its release in the supernatant was largely reduced in the VDU^GFP^ (Figure 7F and G). We then measured the TS activity, as the mAb39 antibody recognizes both active forms of TS and inactive TS, in which Tyr342 is replaced by His342 [64]. The VDU overexpressor and, to a lesser extent the VDU-KO cells, presented a reduced TS activity in the supernatant, matching the decreased TS enzyme levels of the secreted enzyme as detected by mAb39 reactivity (Figure 7G and I). Conversely, TS protein associated with the parasite is significantly decreased in KO-VDU cells only but not in the VDU overexpressor, while the corresponding drop of TS activity is much more pronounced in the overexpressor (Figure 7F and H). Therefore, TcVDU is most like a negative regulator of secreted TS, as TS activity is significantly decreased in the overexpressor. It is tempting to speculate that TcVDU directs the enzyme for degradation and regulates protein traffic to the cell surface.

### Manipulating VDU expression impacts specific proteins in both epimastigote and trypomastigote stages

To better understand the role of TcVDU total parasites lysates of parasites expressing only GFP, VDUGFP, Cas9 and KO-VDU were analyzed qualitatively mass spectrometry. Proteomic analysis identified 5168 proteins in epimastigotes and 4291 in TcT samples (Table S2). TcVDU appears to be at rather low abundance in WT cells, as judged by moderate intensities of detected peptides and the failed detection in a minority of WT replicates. In contrast, overexpressing epimastigote forms showed high levels of the TcVDU peptidase CA C19, with over 500-fold abundance increase (Figure 8A). ER membrane protein 4, involved in protein folding was also augmented. Contrariwise, a subunit of the eukaryotic initiation factor 2B, a guanine-exchange factor for the protein synthesis initiation and a retrotransposon protein were the main proteins with decreased abundance. By analyzing the major protein groups attributed to the different class of proteins, no significant changes in the expression were found (Figure 8B), suggesting that the effect of VDU^GFP^ overexpression may be due to its presence, decreasing the endocytic process and cellular multiplication.

**Figure 8.**
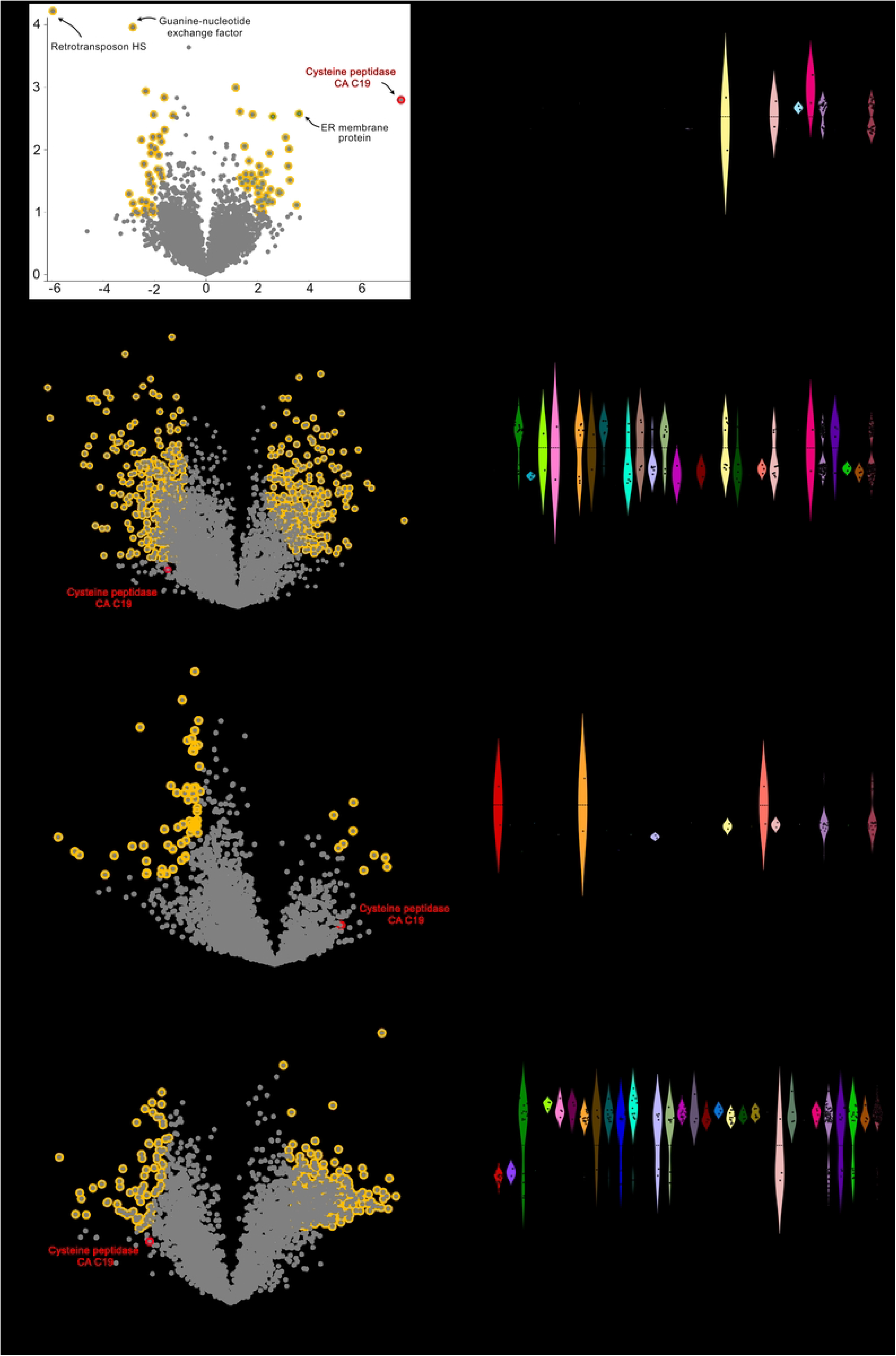
Proteomic analyses of VDU mutants reveal specific reveal protein abundance changes with specific functional enrichments. Duplicate to quintuplicate samples of epimastigote and TcT extracts were subjected to label-free quantification proteomic analysis as detailed in Methods. Volcano plots (t-test difference plotted against the respective -log10 – transformed p values) are shown for protein abundance changes in epimastigotes (A and C) and TcT (E and G) comparing parasites with GFP with VDU^GFP^ and KO-VDU with Cas9GFP. Significant variations are highlighted in orange and selected proteins are annotated. The red dots represent TcVDU. Panels B, D, E and F, show a violin plot of the corresponding differences found for the protein class for genes present in more than 5 haplocopies in the genome. The dots indicate the orange hits and the width, the relative abundance for each class.

A contrasting situation was detected in VDU knockout epimastigote line (Figure 8C). Although some TcVDU expression was detected in the proteomic analysis of epimastigotes KO-VDU (0.03-fold relative to the WT levels), with some corresponding VDU peptides, the presence of additional copies of the gene in some parasites, or incomplete elimination of WT cells in the population may not impact our analysis. In the population, we found increased levels of ribosomal proteins, dyneins, proteins involved in ubiquitination and flagellar attachment zone protein (FAZ) (Figure 8D). At the same time, ABC transporters, protein kinases, proteosome components, glycosyltransferases, proteins involved in protein export were decreased. Of particular relevance is increased expression of SNF-7, a component of the ESCRT system and responsible for final events in endosomal membrane invagination for sorting of ubiquitylated proteins. An increase in SNF-7 may be a response to an increased flux of ubiquitylated proteins resulting from loss of DUB activity. Furthermore, as SNF-7 is the major structural component of ESCRT, it is perhaps to be expected that this protein would be the major subunit to demonstrate altered expression.

Our analyses also showed a contrasting pattern when comparing the overexpressor and knockout of VDU in TcT. In the overexpressing line, the parasites presented several proteins with decreased expression compared to controls (Figure 8E). The level of VDU^GFP^ was increased 9.1-fold in epimastigotes. The increase was less in TcT, probably because the expression was driven by the ribosomal promoter of pTREX, which is less active in this stage. Several conserved hypothetical and unknown proteins decreased (Figure 8F). In the knockout line, we observed the opposite situation, with many groups of proteins increased compared to control parasites (Figure 8G), including many conserved and unknown proteins (Figure 8H). These include many proteins involved in cellular growth, motility, and cell differentiation. In addition to the increased levels of ribosomal and flagellar proteins we observed a large increase in levels of dyneins, mitochondrial proteins, proteases, phosphatases, Rabs, conserved and other identified proteins, suggesting that TcT forms have not shutdown several biological processes as normally found in these stages [65]. Proteasomal proteins, DUBs, and vacuolar proteins mostly augmented. Remarkably, mucin-like glycoproteins and mucin-associated surface proteins (MASPs) largely expressed in the surface of TcT were found diminished in the KO-VDU, but not in the VDU overexpressors.

## Discussion

Changes to the *T. cruzi* surface during the life cycle are key adaptations to the host. *T. cruzi* recognizes signals and nutrients by expressing transporters and receptors at the surface [66–68]. Differentiation into trypomastigotes is accompanied by expression of abundant mucin-like glycoproteins forming a sialylated coat, both protecting the parasite and facilitating infection and survival in the mammalian host [69]. Trypomastigotes also express *trans*-sialidases (TS) crucial to attachment and invasion, and also required for progression to intracellular amastigotes [12]. Remarkably trypomastigotes do not perform endocytosis [27, 50] but are prone to secrete surface proteins. These changes are likely controlled by multiple mechanisms, including alterations to the trafficking machinery.

In this work, we have obtained evidence that disturbing expression levels of the deubiquitinase TcVDU impacts both endocytic membrane trafficking and differentiation in *T. cruzi*. Increasing VDU expression level in proliferating forms of the parasite delayed cargo transfer to late endosomes, whereas the absence favors delivery to the final endocytic compartments. Kinetics and morphological observations by fluorescence and transmission electron microscopy suggested that the action of TcVDU on putative cargo proteins may occur at the late endosome entry point, possibly mediated by direct receptor cargo deubiquitination as shown for the β-adrenergic receptor endocytosis in mammalian cells as well as ISG65 and 75 in the related African trypanosome [70]. This is consistent with TcVDU acting on ubiquitylated proteins, such that over expression decreases ubiquitination levels in endocytosed cargo and promotes recycling while the knockout has the inverse impact. In non-proliferative tissue culture trypomastigotes, overexpressing TcVDU decreased infectivity and the release of active TS. Conversely, decreased TcVDU increased the abundance of several proteins involved in growth, with a decrease of typical surface proteins of TcTs such as MASP, mucins and active TS, suggesting compromised remodeling of the surface. Overall, these data indicate a critical role for TcVDU in both coordinated endocytosis and by consequence the composition of the cell surface, leading to decreased infectivity.

There is considerable evidence that manipulating expression levels of specific DUBs can alter sorting of ubiquitinated proteins and interfere with lysosomal delivery [52], and this is consistent with a requirement for ubiquitylation in delivery of proteins to the reservosome. TcVDU thus has a similar function as mammalian USP33 and USP20, which act to regulate lysosomal delivery [70, 71]. Although the control of endocytosis via TcVDU appears to be conserved in kinetoplastids, only *T. cruzi* perform endocytosis primarily through the cytostome/cytopharynx complex [56], which is distinct from *T. brucei* and *Leishmania spp,* in which receptor mediated endocytosis occurs exclusively via the flagellar pocket [72–75]. Interestingly, TcVDU accumulates at the flagellar pocket, and in both trypanosomes with and without the cytostome, mediates trafficking [76]. Therefore, it is reasonable to propose that VDU in trypanosomes functions predominantly after initial entry and at the sorting endosome when cargo is transferred to subsequent compartments, explaining the abnormal accumulation of transferrin in the cytopharynx and delayed transfer to reservosomes in TcVDU overexpressor cells. In contrast, frequent fusion of endosomes with reservosomes was seen in lines with reduced expression of VDU, consistent with increased endocytic flux. Our evidence that some VDU^GFP^ and V5 tagged-VDU surrounds endosomes could support this assumption, although more detailed studies should be made. In addition, the observed increase in size and reduction in the number of reservosomes in the knockout line further suggest TcVDU role in the membrane traffic, which changes during differentiation to infective trypomastigotes and correlates with the decreased level trypomastigote markers in the knockout, in agreement with observations of decreasing reservosome size upon starvation [77].

Changes in the protein composition of VDU^GFP^ epimastigotes, in comparison with the KO-VDU line, were observed and reveals the importance of the presence of VDU for the control of protein expression in proliferating forms, with 426 proteins decreased and 169 increased at least 2-fold in the KO-VDU. Maximally enriched functional groups in the cohort of significantly increased proteins were dyneins and multiple proteins related to ubiquitination. The increase in dynein levels could be explained by observations that ubiquitination changes affect the dynein localization [78]. DUB1 and 2 regulates dynein levels in B-lymphocytes [79], and at least one DUB (USP21) is directly associated with centrosomes and microtubules in mammalian cells [80]. We detected additionally increased levels of small ubiquitin (C4B63_31g548c), the ubiquitin fold modifier protein (C4B63_35g1396c), increased expression of E2 ubiquitin-ligases (C4B63_32g327/CAB63_340g2), which is structurally similar to UBC12 and 13 ligases and an increase in the HECT domain E3-ubiquitin ligase (C4B63_45g234), known to promote protein proteasomal degradation [81]. In the absence of TcVDU, we also observed an increase in intraflagellar proteins (e.g., intraflagellar transport proteins and flagellar attachment zone proteins), likely explained by the dependence of flagellar assembly and intraflagellar transport on the ubiquitination state of component proteins [82]. Accordingly, mutations in dyneins of *T. brucei* causes significant changes in flagellar formation [83]. SNF7 (C4B63_2g524), a component of the ESCRTIII is also increased, consistent with augmented endocytosis in the KO-VDU, possibly similar to changes in the ubiquitin turnover related to yeast endosomal trafficking [84].

In contrast, we observed decreased levels of Rab 14, 21, 32 and 28 proteins in the KO-VDU in epimastigotes. Rab14 is involved in Golgi to endosome transport and expression of a dominant negative mutant increases the number of secondary lysosomes in mammalian cells [85], similar to the phenotype observed here. In *T. brucei* Rab28 interacts with ESCRT pathways and depletion inhibits endocytosis, consistent with decreased abundance of ESCRTI complex and retromer [86]. Rab21 is important for transfer of primary to secondary endosomes [87] and a decrease in the KO-VDU cells likely related to an analogous role in trypanosomes [88]. Similarly, Rab32 has a unique role in the active exchange of endosomal proteins in *T. cruzi* [89] and *T. brucei* [90]. Significantly, as these changes in Rab GTPases are not seen in VDU1 RNAi in *T. brucei*; we suggest that this is likely the result of different methods used to manipulate VDU in the two parasites. For *T. cruzi* inducible systems are not available so that the parasite has many generations of selection to adapt to changed VDU activity, whereas experiments in *T. brucei* were performed using RNAi, a much more rapid approach and precluding the selection of adapted parasites.

We also investigated the consequences of VDU overexpression and knockout in the mammalian infective stage of the parasite. Decreased infectivity, without changes in the efficiency of egress from infected cells is indicative of an impact on factors related to the invasion process, but not significantly relevant for amastigote intracellular growth and maturation. We observed alterations to the TS expression in both VDU mutants. The KO-VDU exhibits a decrease in the SAPA family, *albeit* modest in magnitude. This includes active enzymes, released from the cell and cell associated enzymatic activity all clearly reduced in the overexpressor line and likely contributing to decreased invasion. It is also important to consider that multiple TS gene families, probably with distinct secretory paths are expressed by the parasite and that VDU likely impacts each of these differentially.

Interestingly, culture-derived trypomastigotes with reduced levels of Tc-VDU showed no clear infectivity phenotype although changes in protein abundance were detected. There was a general decrease in the levels of mucin and MASP, the most abundant proteins of the cultured trypomastigote surface. There is no evidence that these proteins are directly related to cell invasion, and most likely act as a coat that protect the parasite against host immune response, which is unlikely to have any impact in the *in vitro* system used here [91]. In contrast there are no large changes in mucins and MASP in TcVDU overexpressors, but members of each group of the TS gene family are differentially impacted in the KO-VDU (Figure S8). Group V, the most abundant are equally distributed, while group II, represented by glycoproteins involved in host cell attachment have some members with increased expression in the KO-VDU [59], while others are decreased. The majority of group VI and VIII proteins have decreased expression in the TcT KO-VDU. It must be born in mind that the impact of TcVDU on mucins, MASP and TS is likely in part secondary, as the DUB can only act directly in *trans-*membrane domain proteins and not lipid-anchored ones.

Trypomastigotes (metacyclics and TcT) are parasite stages that synthesize mainly surface proteins relevant for infectivity and host interaction [92–94], which are continuously released by the parasite [29, 30]. Therefore, the observed increased levels of many proteins involved in cellular growth occurred in TcT KO-VDU suggests that the released parasites are still more metabolically active, and not completely differentiated compared to wild type TcT. Based on the observed reversion of traffic upon VDU depletion, which in fact increased the formation of larger secondary endosomes or reservosomes, it is tempting to speculate that transition from typical endocytic character to a secretory parasite requires VDU. It is then possible that these incompletely differentiated cells can invade cells as has been demonstrated recently for recently differentiated epimastigotes [95].

A distinct impact of VDU change has been observed in *T. brucei* depending on the mode of membrane attachment. Proteome changes observed upon RNAi depletion of the VDU ortholog in *T. brucei* are different [32]. The impact of TbVDU *knockdown* is limited to a defined cohort of *T. brucei* specific surface proteins, namely ISG65, ISG75 and ISG-related, while the whole cell proteome is not significantly perturbed. Notably, as RNAi does not result in complete depletion, it is possible that the residual pool of TbVDU is sufficient to maintain various other cellular functions. Notably, *T. brucei* VDU localizes to the flagellar pocket and early endocytic compartments, where it appears to control trafficking of ISGs that are frequently internalized by endocytosis [32], which bears some resemblance to the role in trafficking of VDU1in *T. cruzi*.

In conclusion, a conserved deubiquitinase is relevant for control of endocytic traffic in *T. cruzi*. Deubiquitination by TcVDU appears to prevent transfer of cargo from the cytopharynx to endosomes, a process employed for nutrient scavenging in insect stages of the parasite. Importantly, in non-proliferative and infective parasites, the deubiquitinase controls the surface composition, and which is required for efficient mammalian host cell invasion.

## Material And Methods

### Ethics statement

All experiments of this work, including Animal welfare and ethical review board were performed were approved by the Ethics commission of the Federal University of São Paulo from Brazil, under the number CEUA 266629105. The use of recombinants was evaluated and approved by the Internal Commission of Biosafety (CiBio) from the Federal University of São Paulo (2016/3) followed by certification by the Brazilian National Committee of Biosafety (CTNBio).

### Parasite and mammalian cell cultures

*Trypanosoma cruzi* epimastigotes (Dm28c strain) were maintained in exponential growth phase in liver and infusion tryptose medium (LIT) [96] supplemented with 10% fetal bovine serum (FBS) and 10 U/ml penicillin/10 μg/ml streptomycin (Invitrogen) at 28°C. Parasites were usually seeded at 5 x 10^6^ epimastigotes/mL (logarithmic growth phase) and counted manually in a Neubauer chamber or with MUSE equipment when performing growth curves (Thermo Fisher Scientific). For differentiation, epimastigotes were grown until they reached a stationary phase (5×10^7^ cells/ml), and then the differentiation was accomplished as previously described [97]. Metacyclic trypomastigotes were purified from the culture supernatant by ion-exchange chromatography using 2-(diethylamino) ethyl ether cellulose (DEAE-cellulose) columns [98]. *T. cruzi* tissue-culture derived trypomastigotes (TcT) were obtained from supernatants of LLC-MK2 cells (Rhesus monkey kidney epithelial cells, ATCC CCL-7), or U-2OS cells (Human osteosarcoma cells, Banco de Células do Rio de Janeiro) maintained as described [99]. The cells were infected with metacyclic-trypomastigotes, or with trypomastigotes released from infected cells. The parasites were collected by centrifugation at 1500 g for 5 min, incubated for at least 60 min at 37°C and trypomastigote enriched supernatants collected.

### Invasion, intracellular growth assays

For infectivity assays, U2OS cells (4000 cells/well) were seeded in 96 well black plates with clear bottom in 0.1 mL of low glucose DMEM media supplemented with 10% fetal bovine serum (FBS) and incubated for 24 h at 37°C and 5% CO_2_. The cells were then incubated with tissue-culture trypomastigotes (TcTs) from U-2OS cells at a multiplicity of infection (MOI) 10 in 100 µL per well. For analysis of infection ratio, non-invading parasites were removed after 24 h, wells washed with PBS, and the cells were fixed with 4% paraformaldehyde (PFA) in PBS and stained with Draq5 (Biostatus). The percentage of infected cells and the mean parasite number per cell was determined by imaging with High Content Analysis System INCell (GE), with a 20x magnification.

To monitor the intracellular growth, the MOI was adjusted to obtain the same infectivity for each parasite line. The parasites were incubated for 24 h with the cells, the wells were PBS washed and incubated with fresh medium for further 48 h hours. The number of intracellular amastigotes were quantified as above as described previously [99]. In parallel, TcT egress was determined by counting the number of parasites released in the culture supernatant.

### Plasmid construction

PCR fragments containing the *T. cruzi* VDU gene (BCY84_01319, http://tritrypdb.org), and USP-7 gene (<u>BCY84_13892</u>) homologues respectively to the *Trypanosoma brucei* VDU Tb927.11.12240 and USP7 <u>Tb927.9.14470</u> [32] were amplified from extracted DM28c epimastigotes genomic DNA, extracted as in [100] by using the primers TcVduXbaIFow and TcVduBamHIRev / Usp7XbaFow and Tcusp7BamRev. The resulting amplicons were digested with XbaI/BamHI and inserted into pTREX-GFPSBNeo [44] pre-digested with XbaI and BamHI. Primer sequences are given in Table S1. The final constructs were termed pTREX-VDUGFP and pTREX-USP7 and used to transfect exponentially growing epimastigotes. In parallel parasites were transfected with pTREX-GFP-Neo or pTREX-CAS9-GFP-Neo [49]. Briefly, 1 x 10^7^ epimastigotes in 0.5 mL were incubated with 30 μg of the plasmids in the buffer as described [101] and pulsed twice with the AMAXA Nucleofactor II electroporator (Lonza) X-014 program. The *T. cruzi* transformed lines were selected in LIT medium supplemented with 10% FBS and 200 μg/mL G418 and the presence of GFP expression was confirmed by flow-cytometry and fluorescence imaging.

### TcVDU knockouts

For the generation of VDU knockout parasites and V5-VDU tag, Cas9 protospacer adjacent motif (PAMs) with 20 bp protospacers were screened based on the *T. cruzi* VDU coding sequencing using the EuPATGDT tool. Based on the most adequate regions, we selected the oligonucleotide TcVdu-Primer sgRNA184-FOW for the knockout and sGVDUTag for the V5 tag, designed to include the T7 RNA polymerase promoter (italic); the protospacer region of 20 nucleotides (bold) followed by a region of 23 bases that anneals to the pUC-sgRNA template plasmid [49]. These primers together with the scaffold sgRNAREV primer were used to amplify a fragment of the sgRNA template (82 bp) using pUC-sgRNA as template. For the knockout, the PCR product was used as template for *in vitro* transcription with MEGAscript™ T7 Transcription Kit (Thermo Fisher Scientific) as described by the manufacturer to generate sgRNA184 RNA. For the V5 tag, the PCR product was directly used for transfection. The donor sequence encoding blasticidin (BSD) resistance flanked by the 5’ region of VDU and part of the VDU coding sequence was prepared by PCR using the plasmid pTREX-b-NLS-hSpCas9 (a gift from Rick Tarleton, Addgene plasmid #62543) as template and the donor TcVDU84BlastFOW and donor TcVDU84BlastREV. The sgRNA (2 µg) and the donor DNA (25 µg) were used to transfect CAS9::GFP epimastigotes as described above.

To generate the V5 tag, the donors were generated by PCR reaction using as template the total *T. cruzi* DNA with P1VDUv2 and V5TagRev. The product was reamplified with P1VDUv2 and VduV5Rev and the product together with the sgRNA template used to transfect epimastigotes containing pLEW-T7Cas9-Neo plasmid linearized with *Not* I, for expression of Cas9 and T7 RNA (Dm28c^T7 Cas9^) [102]. In both cases, parasites were selected in LIT medium supplemented with 50 μg/mL of BSD and 100 μg/ml of G418. The deletion of TcVDU was verified by PCR of the parasite DNA with the primers TcVDU_KO2_FOW (P1), TcVDU_KO2_REV (P2), TcVDUNtRTF (P5), TcVduRTRev (P6), TcVDUF1 (P7), and TcVDUKO184Rev (P8). The following primers were used to confirm the insertion of the BSD gene (BLASTKOfw, P3, and BLASTKOrev, P4). The PCR products were examined by standard agarose gel electrophoresis [103].

### qPCR analysis

Total RNA was isolated from exponentially growing epimastigote (1 X 10^8^) parasites using Trizol (Thermo Fisher Scientific) as recommended and treated with 2 U of Turbo DNAse for 30 min at 37°C. The resulting samples were further extracted with acid phenol, washed with CHCl_3_ and ethanol precipitated and resuspended in RNase free water. 10 µg of RNA was denatured in the presence of 10 pmoles of oligo dT, 1 mM dNTPs at 65°C for 5 min and annealed for 30 min on ice. cDNA was prepared with 2 µg of denatured and annealed RNA by incubation for 50 min at 50°C in reverse transcriptase buffer, 2.5 mM MgCl_2_, 40 U of RNase out and 200 U of Superscript III (Thermo Fisher Scientific). The reactions were terminated by 5 min incubation at 85°C and treatment with RNAse H for 20 min at 37°C. PCR amplifications were performed in a Bioer FQD96A thermocycler with 0.04 µg of cDNA, 2.5 nmol of each primer and Sy after 10 min at 94°C with 40 cycles (15 sec 94°C, 20 sec 60°C) and standard melting curves. For the N-terminal portion of VDU we employed primers VDUNtFor and VDUNtRev and for the C-terminal portion the VDURTFor and VDURTRev. The expression levels were calculated by the -ΔΔCt method using primers for *T. cruzi* GAPDH (GAPDHF and GAPDHR), all listed in Table S1.

### Western blotting

Whole parasite extracts were prepared by collecting cells and boiling in 5x Sample Buffer (250 mM Tris. HCl, pH 6.8, 10% SDS, 30% (v/v) Glycerol, 10 mM DTT, 0.05% (w/v) Bromophenol Blue). Extracts were resolved on 10% polyacrylamide gels and transferred to nitrocellulose membranes (GE Life Sciences) by standard procedures. The membranes were stained with 0.3% Ponceau S (w/v) in 0.3% (v/v) acetic acid, washed in water and blocked with 5% (w/v) non-fat dry milk in 10 mM Tris-HCl, pH 7.4, 0.15 M NaCl and 0.05% Tween-20 (TBS-T). Primary antibody incubation was performed for 1 h at room temperature with TBS-T supplemented with 5% non-fat dry milk. Membranes were washed with TBS-T and then incubated with secondary antibodies in the same conditions as the primary antibodies. IRDye-conjugated secondary antibodies (anti-rabbit or anti-mouse IgG coupled to IRDye 800 and IRDye 680 LI-COR Biosciences) were used for protein detection and visualization. Proteins were detected by western blotting using the following antibodies: Anti-GFP antibody (1:3000 dilution; Millipore), anti-aldolase antibody (1:3000) [104] and mAb39 anti-trans-sialidase antibody (1:1000) [105].

### Transferrin endocytosis

Log-phase epimastigotes (2 x 10^7^ /mL) were washed in RPMI 1640 (Thermo Fisher) without FSB, and then incubated in 100 μL of the same medium for 15 min at 28°C. Transferrin from human serum, conjugated to Alexa Fluor 633 (Thermo Fisher Scientific) was added to a final concentration of 50 μg/mL. Cells were incubated for 1, 10, and 30 min at 28°C and then immediately fixed in 4% PFA in PBS for 20 min and examined by fluorescence microscopy and flow cytometry. Dynamics of transferrin endocytosis over time (min) was evaluated by manual counting of positive cells to transferrin labeling in fluorescence microscopy images. One hundred cells were counted per assay in each condition. The endocytosis was also measured by flow cytometry by using a BD Accury C6 apparatus (BD Biosciences) by considering the transferrin-Alexa 633 (TF^633^) fluorescence, which was measured in FL4-A channel (660/20 filter). The quantitative analysis was made calculating the percentages of fluorescent cells and the total incorporation, in both cases by subtracting the fluorescence intensity of parasites incubated without TF^633^ from the fluorescent of parasites incubated with TF^633^. Cell cycle profile was determined in parasites (1 x 10^6^/mL) fixed with 70% ethanol, washed, resuspended in PBS, and incubated with 0.25 µg/mL 7-Aminoactinomycin D (7-AAD) for 10 min. The parasites were collected by centrifugation, washed once with PBS, and analyzed by flow cytometry in the FL40A channel. Data were analyzed with FlowJo software (BD Biosciences). Negative controls, which included cells incubated in the absence of transferrin or cells without GFP expression, were used to select gates to discriminate positive and negative events and 20,000 events were analyzed per sample. Fluorescence images were acquired using an Orca R2 CCD camera (Hamamatsu) coupled to an Olympus BX-61 microscope and a ×100 plan Apo-oil objective (NA 1.4). Z-series were acquired from each field using the Cell^M software (Olympus) and processed for 3D-deconvolution using the Autoquant 2.2 software (Media Cybernetics).

### Electron microscopy

For transmission electron microscope (TEM), epimastigotes previously washed in PBS and resuspended at 1 x 10^7^ cells/ml were incubated for 30 min at 28oC in RPMI 1640 without FBS, containing 100 µg/mL of bovine holotransferrin (Sigma-Aldrich) coupled to gold particles (8-10 nm), as described previously [106]. The parasites were then fixed with glutaraldehyde added directly to the incubation medium (2.5% final concentration) and processed as described [56]. Further, the cells were post-fixed using an osmium-thiocarbohydrazide-osmium (OTO) protocol [56, 107], dehydrated in acetone and embedded in epoxy resin. Ultrathin sections were collected in copper grids, post-embedded stained with 5% uranyl acetate and lead citrate and observed in a Tecnai Spirit (Fei Company) microscope operating at 120 kV.

### Focused Ion Beam-Scanning Electron Microscopy (FIB-SEM) and three-dimensional reconstruction

Samples processed for TEM were imaged using FIB-SEM. For this, embedded samples were trimmed to a trapezium shape in a Leica UC7 ultramicrotome and the top of the block was glued to a SEM stub using carbon tape and metalized with gold. Samples were imaged using an Auriga dual-beam microscope (Zeiss) equipped with a gallium-ion source for focused-ion-beam milling, a field-emission gun, and an in-lens secondary electron detector, for SEM imaging. The cross-sectional cut was made at ion beam currents of 2.0 µA and an accelerating voltage of 30 kV. Back-scattered electron images were recorded at an accelerating voltage of 1.8 kV and a beam current of 0.8 nA, in the immersion lens mode, using a CBS (Concentric Back Scatter) detector. A series of backscattered electron images were recorded in ‘slice-and-view’ mode, at a magnification of 15K, with a pixel size of 5.8 nm and milling step size of 30 nm. The sequence of images acquired were aligned, and the volume was reconstructed, allowing stereological analyses of the organelles using IMOD software [108].

### Trans-sialidase assay

Parasites were previously washed in PBS and resuspended to 2 x 10^7^/mL of low glucose DMEM medium containing 10% FBS and incubated for 2 h at 37oC. The cells were collected by centrifugation (2000 g, 5 min) and lysed in 50 µL of 50 mM Tris HCl, 0.1 M NaCl, 1% Triton-X100 and Complete C, EDTA-free protease inhibitor from Roche Diagnostics at 4°C. Different volumes of lysates and the remaining supernatant were incubated 30 min at room temperature in 50 µL of 20 mM Hepes buffer pH 7.0, 1 mM sialyl-lactose (Sigma) containing 0.2% BSA, and 7.2 µM of [D-glucose-^14^C] lactose (60 mCi/mmol, New England Nuclear) [109]. The mixture was diluted in 1 mL of water and added to a QAE-Sephadex-A50 column (1 mL equilibrated in water). After washes with 8 mL water, the labeled sialyl-lactose was eluted with 1 M ammonium formate and quantified by scintillation counting using a Tri-Carb 2810 TR counter (Perkin Elmer).

### Proteomic analysis

Epimastigotes and trypomastigotes lysates containing 2 x 107 parasites in 20 µL of SDS-PAGE sample buffer containing 5 mM DTT were boiled for 5 min and loaded onto precast Novex Value 4-12% Tris-Glycine gels (Thermo Fisher). The electrophoresis was performed in 50 mM MOPS-50 mM Tris Base, 3.5 mM SDS, 1 mM EDTA at 100 V for 10 min. The migration portion of each lane were cut, and the gel pieces were fixed for 10 min at room temperature in 1 volume of 40% ethanol, 10% acetic acid. The fixative was removed, and the fixation repeated twice more. The bands were kept frozen at -80oC until processing for mass spectrometry analysis. The gel slices were subjected to reductive alkylation and in gel tryptic digest using routine procedures. The eluted peptides were then analyzed by liquid chromatography-tandem mass spectrometry (LC-MS2) on a ultimate3000 nano rapid separation LC system (Dionex) coupled to a Q Exactive HF Hybrid Quadrupole-Orbitrap mass spectrometer (Thermo-Fisher Scientific). Spectra were processed using the intensity-based label-free quantification (LFQ) method of MaxQuant [110] searching the T. cruzi DM28c-2018 annotated protein database from TriTrypDB [111]. Minimum peptide length was set at six amino acids, isoleucine and leucine were considered indistinguishable and false discovery rates (FDR) of 0.01 were calculated at the levels of peptides, proteins and modification sites based on the number of hits against the reversed sequence database. When the identified peptide sequence set of one protein contained the peptide set of another protein, these two proteins were assigned to the same protein group. The LFQ data were analyzed using Perseus software [112]. A student’s t-test comparing the control sample group to the respective modified strain generated -log10 t-test P-value plotted versus t-test difference to generate a volcano plot. Enrichment analysis was performed using a Fisher’s exact test in Perseus with a Benjamin–Hochberg false discovery rate of 0.02. Proteomic data was also analyzed and visualized using the Orange3 Data Mining (https://orange.biolab.si). Proteomics data have been deposited to the ProteomeXchange Consortium via the PRIDE partner repository [113] with the dataset identifier PXD031835.

### Other statistical analysis

The gene and amino acids sequences of TcVDU of *T. cruzi* were obtained from the TriTrypDB [111], and HsVDU (Q8TEY7) and HsUSP3 (Q9Y6I4) of *Homo sapiens* were obtained from UniProt (https://www.uniprot.org/). To analyze the primary sequence of TcVDU we used the following programs: Predict protein molecular weight and amino acid composition, analysis by (https://web.expasy.org/translate/), Motif analysis by PFAM database (https://pfam.xfam.org/) and multiple sequence alignments were created in Jalview [114] by using the Clustal Omega package [115]. Phylogenetic trees were generated Alignment using MAFFT [116] with the ‘linsi’ option for greater accuracy. Trimming was performed using TrimAl [117] with a gap threshold of 0.7. The substitution model used for both PhyML and MrBayes tree searches was the LG matrix [118], with four gamma-distributed rate categories (LG+G4). Other statistical analyses were performed using Prism 9.0 software (GraphPad, San Diego, CA). Values of p < 0.05 were considered significant using one-way ANOVA used to compare more than two population groups. Acknowledgments We thank Claudio Rogerio Oliveira and Claudeci Medeiros for technical support. We also thank to the technical assistance of Dr. Lorian Cobra (Centro Nacional de Bioimagem, Rio de Janeiro, Brazil) for the acquisition of FIB-SEM images and Lael Barlow for help with the phylogenetic analysis.

## Author contributions

Conceptualization: S.S, M.C.F., N.S.M.; Methodology: N.S.M, N.B.S, S.S. G.P.S., L.M.A. C.L.A. M.Z., N.L.C.S., M.Z.; Formal analysis: M.A.F., S.S., M.Z. N.L.C. N.S.M; Investigation: N.S.M., N.B.S, G.P.S., A.M.M., L.M.A. C.L.A. M.Z.; Data curation: S.S., M.Z. M.A.F. N.L.C. ; Writing – original draft: N.S.M., S.S.; Writing – review & editing: N.S.M., S.S., M.A.F, M.Z., L.M.A., C.L.A., N.L.C.S.; Visualization: N.S.M., M.Z., N.L.C.S., C.L.A.; Supervision: S.S., M.C.F, N.L.C.S.; Project administration: S.S.; Funding acquisition: S.S., M.C.F., M.Z., N.L.C.S.

## Data availability

The mass spectrometry proteomics data have been deposited in the ProteomeXchange Consortium via the PRIDE partner repository with the dataset identifier PXD031835.

## Supporting information

**Figure S1. VDU^GFP^ and USP7^GFP^ are expressed at similar levels in *T. cruzi* epimastigotes**. (A) Western blot of epimastigote extracts of WT DM28c (WT), VDU^GFP^, VDU^GFP^ revealed with anti-GFP antibodies. (B) The respective membrane previously stained with Ponceau S. (C) Flow cytometry analysis of the same set of epimastigotes.

**Figure S2. Expression of VDU^GFP^ in intracellular amastigotes.** Fluorescence microscopy of intracellular amastigotes (Ama) expressing the GFP alone or VDU^GFP^. Panels show the GFP, DAPI and merged fluorescence. N: Parasite nucleus. K: Parasite kinetoplast. Nc: mammalian cell nucleus. Bars = 2 um.

**Figure S3. Localization of VDU tagged with V5 epitope at the N-terminus.** T7-Cas9 epimastigote (A) or T7-Cas9 epimastigote transfected with the BSD-V5 donor selected for BSD resistance (B) were fixed, permeabilized and probed with anti-V5 antibodies followed by anti-mouse IgG-Alexa488 and DAPI. The panels show the corresponding DIC images and sections obtained after 3D deconvolution of DAPI, Alexa488 and merged channels. Arrows indicate the localization of V5 tagged VDU in regions near the flagellar pocket. Bars = 2 mm.

**Figure S4: VDU disruption mutants are viable, whereas VDUGFP overexpression leads to a growth phenotype in epimastigotes.** (A) CRISPR/CAS9 strategy was used to disrupt the VDU gene by the insertion of BSD resistance gene in parasites expressing Cas9. The black arrowhead above VDU gene (blue box) represents the region targeted by the sgRNA and black arrows indicate the position of the primers used for confirmation of the insertion by PCR analysis in the wild type (top) and mutated parasites (bottom). Agarose gel electrophoresis of PCR products obtained with total DNA of the indicated lines with the P1/ P2 primer set (B), with P3/P4 primers set (C), and with P5/P6 and P7/P2 primers sets (D) showing the position of the of 979 bp fragment in WT and Cas9 parasites and of the 1174 bp in the KO-VDU, the resistance gene amplification in KO-VDU and the BSD plasmid with the fragment of 399 bp, the absence of the 566 bp in the KO-VDU and amplification of the remaining portion of the endogenous gene with 316 bp in both lines, respectively. (E) RT-qPCR analysis of the indicated samples using GAPDH as endogenous control and the N- and C-terminus pair of primers of VDU. The numbers are means of 3 independent samples, each measured in triplicate.

**Figure S5. Endocytosis is not affected by USP-7 overexpression**. (A) Epimastigotes transfected with pTREX plasmid containing VDU^GFP^, or USP7^GFP^ were incubated with TF^633^ endocytosis in RPMI medium for 2, 10 and 30 min at 28°C, fixed in 4% and analyzed by flow cytometry. (B) The median fluorescence of four independent experiments shown the relative fluorescence compared to parasites incubated without TF^633^.

**Figure S6. Some transferrin endocytic particles colocalize with the VDU^GFP^.** Endocytosis experiments were performed by incubation of parasites in RPMI medium with TF^633^ for 10 min at 28°C. The parasites were then fixed for 20 min with 4% p-formaldehyde in PBS, washed in PBS and attached to poly-L-lysine coated glass slides. The slides were then treated with DAPI and imaged by a fluorescence microscope. The images correspond to one Z-section generated after blind deconvolution, showing the VDU^GFP^ (green), TF^633^ (red), and DAPI (blue). White arrowheads indicate the superposition of TF^633^ and VDU^GFP^. Bars = 5 µm.

**Figure S7. Colocalization analysis of VDUGFP and TR^633^.** (A) Histogram of the normalized frequency of cells versus the bright detail similarity of the localization of green fluorescence of GFP compared to the TR^633^ red fluorescence of epimastigotes incubated 10 min with TR^633^ (25 µg/mL) generated by the IDEAS software of Luminex from 12160 single epimastigote images. We estimated that higher bright signal was above 2 showing that in about 4.68 % of cells the red coincided with the green dots. (B) Selected images with high coincident signals of the indicated fluorescence. Bars = 5 µm.

**Figure S8. Comparative expression in TcTs of proteins coded by the trans-sialidase gene family in VDU-KO**. Volcano plot of t-test difference plotted against the respective – log10 X transformed *p* values for abundance of the indicated trans-sialidase groups in KO-VDU and control parasites. The data was extracted from Table S2.

**Video V1. 3D analysis of TR^633^ and VDU^GFP^.** Autoquant 2.2 three-dimensional image of a VDU^GFP^ epimastigote after 10 min incubation with TR^633^.

**Table S1.**
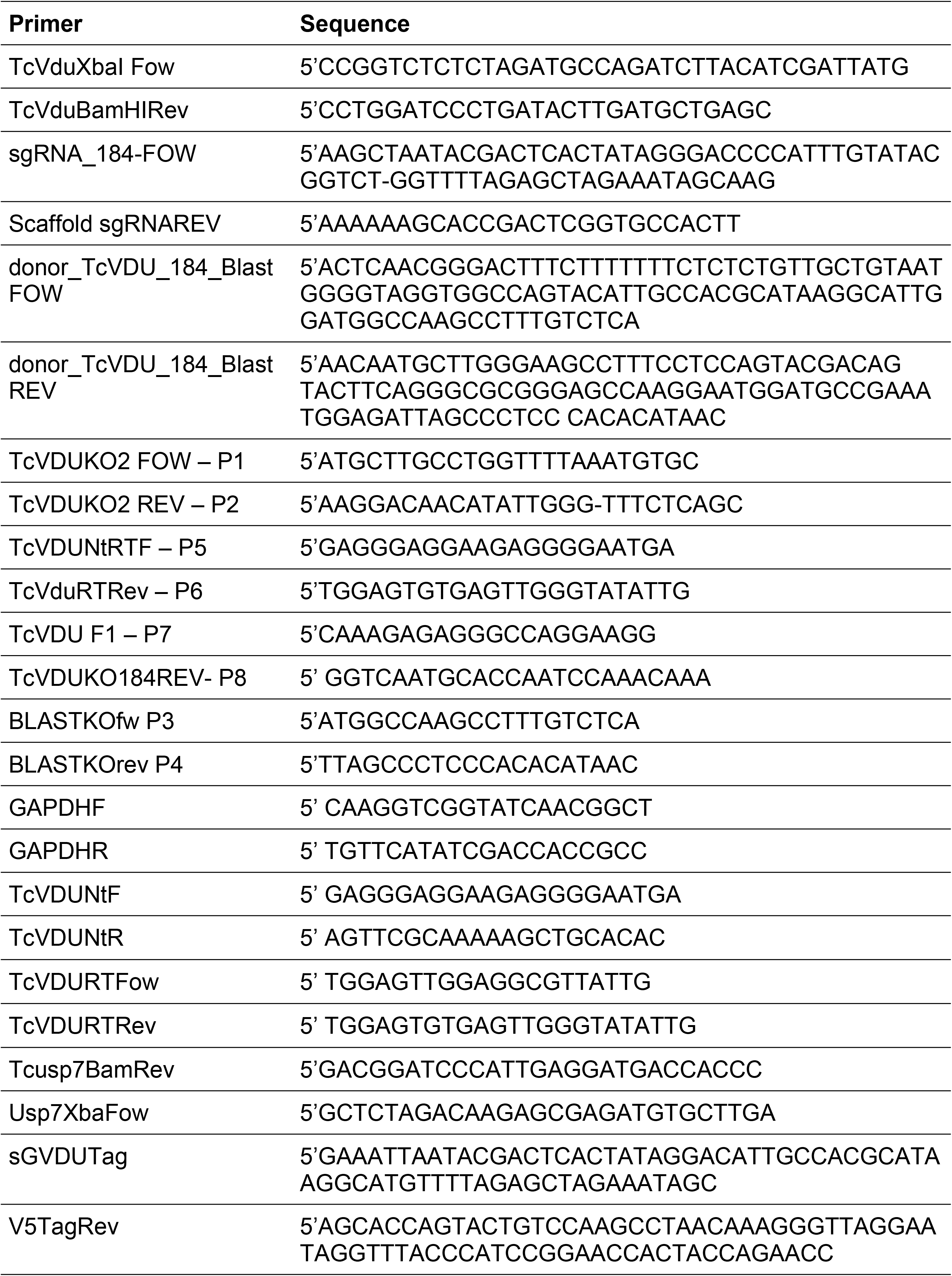
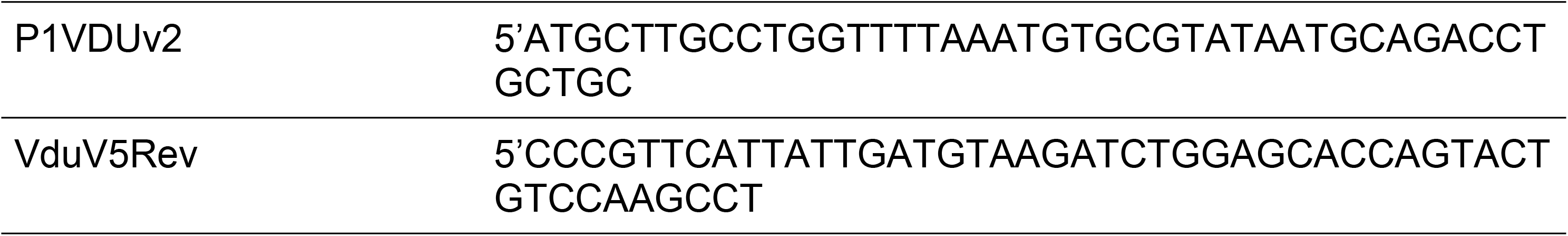
Primers used in this work.

**Table S2. Proteomic data and statistical analysis (see excel file: TableS2.xlsx)**

Tables listing all protein groups quantified for TcVDU overexpression and knockout in epimastigotes (sheets EpiOE and EpiKO) and trypomastigotes (sheets TcTOE and TcTKO) and including corresponding LFQ-intensities, protein annotation, ratios (infinite ratios, t-test difference, -log10 p-value, significance intervals (class A: FDR=0.01, s0=0.1; class B: FDR=0.05, s0=0.1), relevant global search file data, transmembrane prediction (# TM Domains) and GO terms. The two ‘summary’ sheets report sample organization and peak detection statistics for the analysis of epimastigotes and trypomastigotes samples, respectively. In principal component analysis (PCA), the VDU^GFP^ biological replicates were compared to the Dm^GFP^ control and KO-VDU to DmCas9^GFP^.

All raw data (spectra, Maxquant search files and search databases) are available at the PRIDE partner repository [113] under dataset identifier PXD031835.

**Video V1. Similar localization of TR^633^ and VDU^GFP^.** Autoquant 2.2 three-dimensional image of a VDU^GFP^ epimastigote after 10 min incubation with TR^633^.

### Submission details for PRIDE (not for publication)

Project Name: Trypanosoma cruzi VDU deubiquitinase mediates surface protein trafficking and infectivity

Project accession: PXD031835

Project DOI: Not applicable

Reviewer account details:

Username

Password: f7907aRJ

